# All-optical interrogation of a direction selective retinal circuit by holographic wave front shaping

**DOI:** 10.1101/513192

**Authors:** G.L.B Spampinato, E. Ronzitti, V. Zampini, U. Ferrari, F. Trapani, H. Khabou, D. Dalkara, S. Picaud, E. Papagiakoumou, O. Marre, V. Emiliani

## Abstract

Direction selective (DS) ganglion cells (GC) in the retina maintain their tuning across a broad range of light levels. Yet very different circuits can shape their responses from bright to dim light, and their respective contributions are difficult to tease apart. In particular, the contribution of the rod bipolar cell (RBC) primary pathway, a key player in dim light, is unclear. To understand its contribution to DSGC response, we designed an all-optical approach allowing precise manipulation of single retinal neurons. Our system activates single cells in the bipolar cell (BC) layer by two-photon (2P) temporally focused holographic illumination, while recording the activity in the ganglion cell layer by 2P Ca^2^ imaging. By doing so, we demonstrate that RBCs provide an asymmetric input to DSGCs, suggesting they contribute to their direction selectivity. Our results suggest that every circuit providing an input to direction selective cells can generate direction selectivity by itself. This hints at a general principle to achieve robust selectivity in sensory areas.

## Introduction

A major goal in neuroscience is to understand the circuits that embody the computations performed by sensory neurons. In particular, a striking feature of sensory processing is the ability of neurons to perform the same computation in different contexts. For example, neurons in the piriform cortex represent odor identity while remaining invariant to its exact concentration (Bolding and Franks, 2018). Similarly, neurons in the visual cortex can keep the same orientation tuning curve over different contrasts (Sclar and Freeman, 1982).

In the retina, direction selective (DS) ganglion cells (GC) respond selectively to a motion direction in different contexts such as different backgrounds (Chen et al., 2016), different natural scenes (Im and Fried, 2016), and over a broad range of luminosities (Pearson and Kerschensteiner, 2015; Vaney et al., 2001; Yao et al., 2018). Switching the average luminance from dim to bright light will change the dominant circuits that convey visual information inside the retina, leading to major changes in the responses of ganglion cells (Pearson and Kerschensteiner, 2015; Wässle, 2004). For example, cells will change their ON-OFF polarity (Tikidji-Hamburyan et al., 2015). It is therefore particularly striking that direction selectivity remains constant across the ten billion fold change of luminance separating day from night (Yao et al., 2018), despite the involvement of distinct circuits. A major challenge is to understand how this feature is achieved.

During daylight conditions, where rods are saturated and cones transmit light input, the major component responsible for direction selectivity is a functional asymmetry. This results either from an asymmetric inhibition or an asymmetric morphology. For example, in the ON-OFF DS circuit, cones activate GCs via the cone bipolar cells (CBC), but they also activate starburst amacrine cells (SAC) (Kim et al., 2014), which provide a direction selective, asymmetric inhibitory input to DSGCs (Fried et al., 2002; Vaney et al., 2012; Wei et al., 2011) (Fig. 1A). In other DS types, the asymmetric morphology of the dendritic arbor generates the functional asymmetry responsible for the direction selectivity (Kim et al., 2008; Trenholm et al., 2011)(Fig. 1B).

When light gets dimmer, other circuits come into play (Grimes et al., 2018). Rods are not saturated anymore and transmit their output down to GCs through three different pathways. Two of them, the secondary and ternary pathways, are only one synapse away from the cone circuitry. In the secondary pathway, rods directly communicate with cones through gap junctions (Fig. 1C). In the ternary pathway, rods directly synapse to a specific type of CBC (Fig. 1D). Since rods connect directly to the cone circuitry, they modulate the direction selective circuit and can be used to compute direction selectivity during night vision.

**Figure 1.**
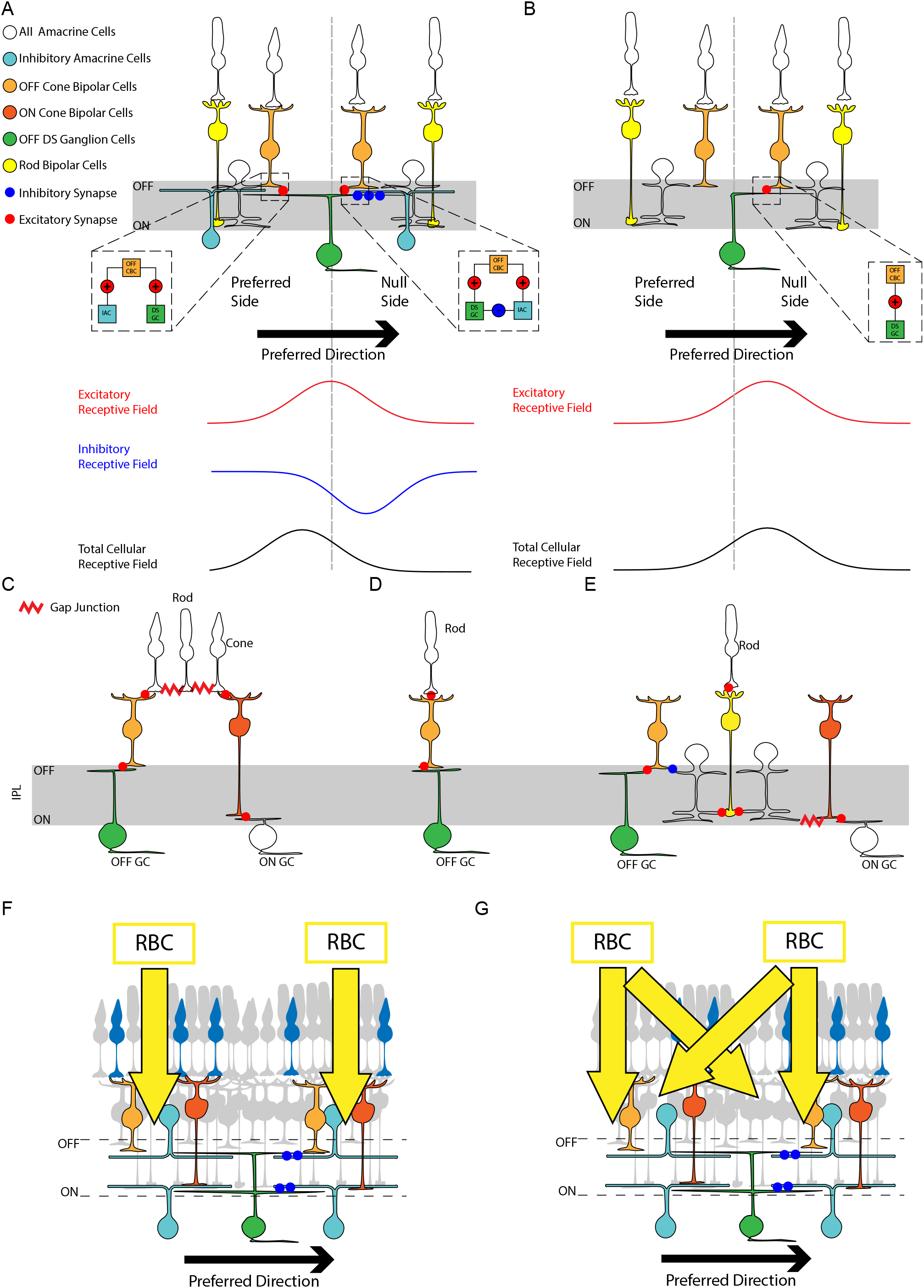
Different circuits to generate direction selectivity give different predictions for the cellular receptive field. **A:** direction selectivity is due to asymmetric inhibition. OFF DSGCs receive symmetric inputs from OFF BPs, responsible for the excitatory receptive field. They also receive asymmetric inhibitory inputs from an AC, specifically from the null side. This input will cancel the excitation when the bar moving from the null side (null direction), but not when moving from the preferred side (preferred direction). This spatially shifted inhibition will also generate a cellular receptive field shifted towards the preferred side. Insets: Schematic description of the circuit at the OFF-CBC synapses with the OFF DSGC responsible for the generation of the direction selectivity on the two sides. On the null side there are synaptic contacts between inhibitory ACs and the OFF DSGC, but not on the preferred side (see inset). **B:** direction selectivity is due to an asymmetric dendritic field. OFF BCs innervate only the null side. This and the non-linear dendritic integration generate a preference for centrifugal (soma to dendrites) motion. In this case, the excitatory receptive field is spatially shifted towards the null side, and so is the cellular receptive field. **C:** Secondary rod pathway: Rod signals are transmitted directly to cones via gap-junctions, then to ON-OFF CBCs and in turn to GCs with corresponding polarity. **D:** Tertiary rod pathway: Rods are directly connected to a specific type of OFF CBC through an ionotropic sign-conserving AMPA glutamate receptor. **E:** Primary rod pathway: in the mammalian retina: rods transfer their signal to RBCs (yellow) through a sign inverting metabotropic glutamate receptor 6 (mGluR6). RBCs relay the received signal via sign-conserving glutamatergic synapses to AII ACs. AII ACs, stratifying in both ON and OFF strata, split the signal to ON CBCs through gap junctions and to OFF CBCs (orange) through sign-inverting glycinergic synapses. CBCs transfer the received signal to the GC layer (ON and OFF GCs, green) following the regular cone circuitry. F: First hypothesis: RBCs preserve the asymmetry and modulate only neighboring CBCs. For example, left RBCs would modulate mostly the CBCs on the preferred side. **G:** Second hypothesis: the RBC pathway is too divergent and, by modulating distant cone BCs, dilutes the functional asymmetry. For example, left RBCs send their input to both the preferred and the null side and similarly for the right RBCs.

On the contrary, in the primary pathway (Fig. 1E), rod signal is transmitted to two or more RBCs, which in turn input to at least 7 AII amacrine cells (AC), each reaching at least 14 CBCs (Tsukamoto and Omi, 2013, 2017). This pathway is therefore strongly divergent and it is unclear if it contributes to shape direction selectivity, or if it delivers an unselective signal to DS cells. In the first hypothesis, RBCs would impact only neighboring CBCs and preserve the asymmetry (Fig. 1F). For example, for a DS cell that prefers left-to-right motion receiving more inhibition from the right, left RBCs would modulate mostly left CBCs, and conversely for the right side (right RBCs reaching mostly right CBCs). In the second case (Fig. 1G), the strong divergence of the RBC pathway impacts even more distant CBCs and dilutes the functional asymmetry. In this case, left RBCs reach both left and right CBCs, and a similar dilution will occur for right RBCs. The functional impact of RBCs onto DSGCs would thus be symmetric and the primary rod pathway would not participate in the computation of direction selectivity, in this scenario.

In summary, either each of the three circuits contributes to direction selectivity with an asymmetric input, or this is only true for the second and third circuit whereas the primary circuit provides an unselective modulation.

To understand which of these two hypotheses prevails it is necessary to manipulate the primary rod pathway, selectively and without damaging the retinal tissue. This requires (i) precise activation of RBCs; (ii) simultaneous recordings from the activated GCs; (iii) the ability to distinguish among the different GCs types.

Existing approaches to study retinal circuits using electrophysiology (Asari and Meister, 2012, 2014) or electrophysiology combined with optogenetics using wide-field illumination (Park et al., 2015) lack either specific targeting of cells or precise stimulation.

To overcome these shortcomings, we designed an all-optical approach enabling activation of one or multiple cells in the BC layer by two-photon (2P) temporally focused holographic illumination (Accanto et al., 2018), while recording the activity in the GC layer by 2P calcium imaging. The system also enables us to perform visual stimulation of photoreceptors to probe the localization of DSGCs. Because the BC and GC layers lay on axially distinct planes (50-70μm apart) the photostimulation path using a two-step phase modulation scheme (Accanto et al., 2018) allows decoupling the photostimulation from the recording plane.

We focused on a specific type of DSGC cells: the OFF DSGC cells (G_2_ cells in (Baden et al., 2016)). For this type, we found that there was a clear asymmetry in the integration of RBC output, with a clear bias towards the preferred side of these cells. This is consistent with our initial hypothesis that the functional asymmetry behind direction selectivity is preserved despite the strong divergence of the RBC pathway. Direction selectivity is maintained at different light levels because the three pathways converging to these GCs all enable direction selectivity individually.

The cutting-edge experimental strategy that we employed here to dissect retinal circuits, opens a broad range of experiments for the investigation of functional connectivity among multi-layered circuits in different brain regions.

## Results

### An all optical system combining holographic front shaping and 2 photon imaging

The optical setup includes three main illumination paths: a 2P temporally focused stimulation via Computer-Generated Holography (CGH), a 2P scanning galvo-based imaging and a visual stimulation via Digital Micromirror Device (DMD)(Fig. 2A). Holographic illumination was provided by a fiber amplifier laser source, emitting at 1030 nm and operating at 500 kHz repetition rate (Satsuma HP, Amplitude Systems, France) enabling efficient 2P optogenetic activation of opsins (Chaigneau et al., 2016)(Ronzitti et al., 2017). Calcium imaging was provided by raster scanning a 920-nm beam delivered by a Ti:Sapphire oscillator (Chamaleon Vision II, Coherent Ltd., USA). Simultaneous spatiotemporally-controlled full-field or moving-bars stimuli were sent to the sample by using a DMD illuminated with visible light (420 nm). The optical system enabled decoupling the imaging and photostimulation planes by multiplexed temporally focused light-shaping (MTF-LS) stimulation (Accanto et al., 2018), allowing an all-optical interrogation of neural circuits laying at axially distinct planes. MTF-LS is based on a two-step process (see Ref. (Accanto et al., 2018) for a detailed description): a first beam-shaper unit (here a static phase-mask) spatially modulates the phase of the incoming illumination beam to produce a holographic 2D shape (here a circular 10-μm diameter spot). The generated intensity pattern is successively focused on a grating to enable temporal focusing, which provides enhanced axial confinement of the illumination pattern at the sample and robust propagating through scattering tissues (Bègue et al., 2013; Papagiakoumou et al., 2013). A further phase modulation provided by a reconfigurable spatial light modulator (SLM) allows dynamically multiplexing the 2D shape laterally and/or axially in the sample volume (Fig. 2A). Here, we demonstrated that the illumination spot could be displaced axially in a range comparable to the distance between GCs and RBCs (i.e., 60 μm-80 μm), while maintaining an axial confinement of the illumination spot below 10 μm (Full Width at Half Maximum-FWHM of the axial intensity distribution) (Fig. 2B,C). This value represents a 2-3 fold improvement with respect to a non-temporally focused holographic spot of the same size and is maintained independently of the number and distribution of spots (Supplementary Fig. 1). We observed nearly uniform FWHMs by positioning 10-μm diameter holographic spots in a field of view (FOV) of 200×200 μm^2^ at 70 μm below the focal plane (FWHM 8.9±0.75 μm, Mean ± Standard Deviation (SD)) (Fig. 2D, Supplementary Fig. 2). The illumination intensity variability was around ±12% (±SD of intensities) across the FOV (Fig. 2E, Supplementary Fig. 2). This indicates that the system can provide nearly homogenous photostimulation of opsin-expressing cells across the RBC layer, while maintaining the imaging focus on the GC layer.

**Figure 2.**
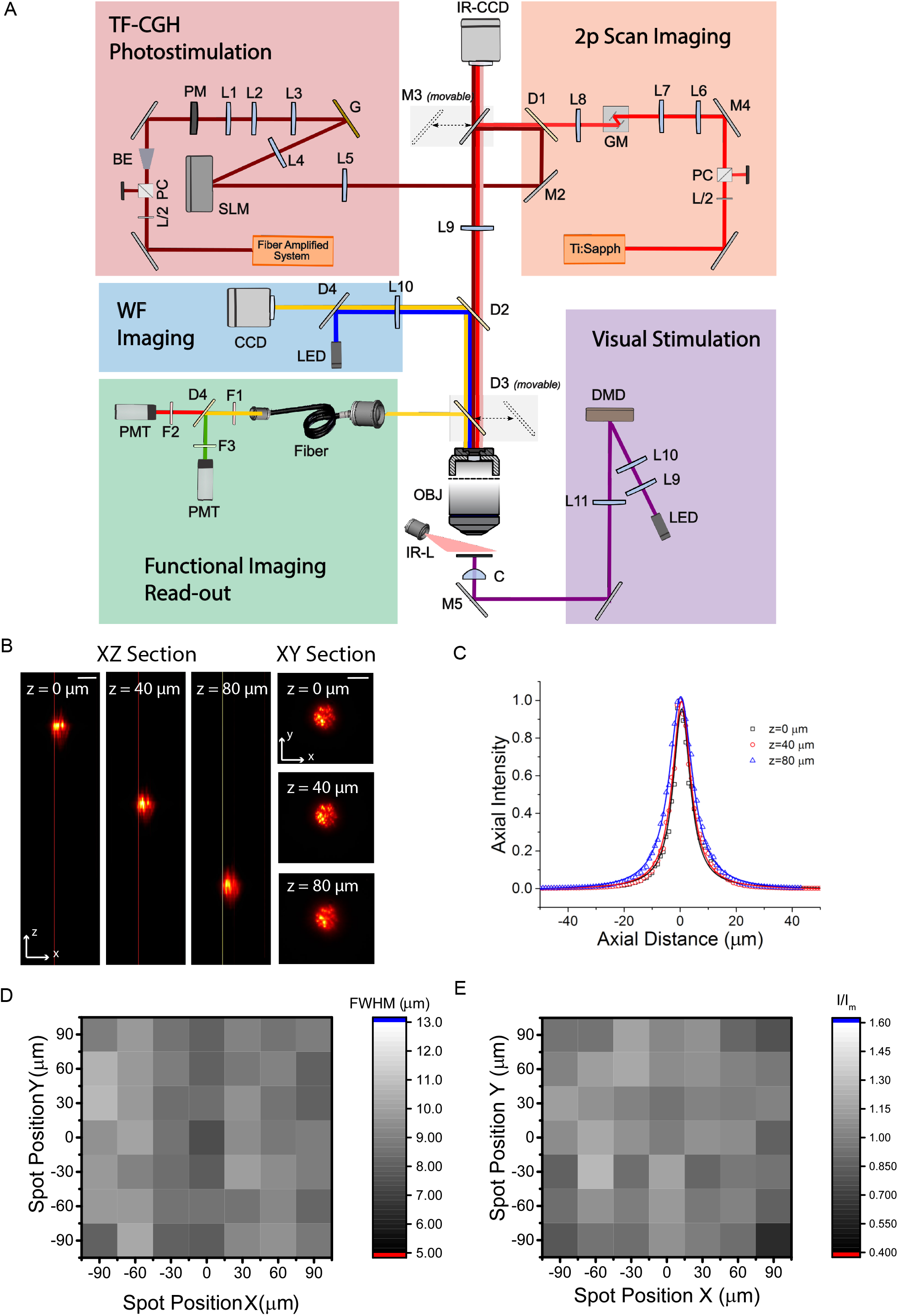
Experimental setup and optical characterization of the system. **A.** The optical system combined a multi-light-path architecture including 2P scan-based, epifluorescence widefield (WF) and infrared (IR) imaging; a holographic-based 3D multiplexing temporally focused photoactivation apparatus for cell activation (TF-CGH Photostimulation) and a DMD-based spatiotemporally-controlled visual stimulation system. **B.** Axial shift of a 10-μm-diameter temporally focused holographic spot. *(Left) xz* orthogonal maximum fluorescence intensity projection of the spot for different axial displacements. *(Right)* Corresponding *xy* 2P fluorescence intensity cross-section. Scale bar, 10 μm. **C.** Axial profile of the fluorescence intensity of spots shown in **B.** Solid lines indicate Lorentzian fit. **D.** 2D map of the Full Width Half Maximum of the axial profiles of the fluorescence intensity given by a matrix of spots (30 μm inter-spots distance) displayed 70 μm below the focal plane (each pixel represents one spot). **E.** Normalized illumination intensity of a matrix of spots displayed 70 μm below the focal plane (each pixel represents a spot).

### Targeted holographic stimulation of RBCs

We injected an AAV in the vitreous of the mouse eye, and expressed CoChR fused with the GFP protein under the control of a promoter specific to RBCs described previously (Lu et al., 2016) (Fig. 3A, see also Methods). To determine the functional resolution of our system we patched RBCs expressing the opsin CoChR, while stimulating them with MTF-CGH. Patching RBCs in a whole mount configuration by going through the GC layer with the pipette is challenging. Instead, we removed the photoreceptor layer and turned the retina upside down to access the RBCs directly (Fig. 3B, C, Methods). This is equivalent to stimulating from the GC side since the retina is a transparent tissue (except for the photoreceptor layer (Chen, 1993)). We patched fluorescent RBCs under 2P guidance (Fig. 3D) and checked the morphology of the patched cell by filling it with Alexa594. We measured the photocurrent under various light intensities which ranged from tens of pA under moderate illumination (32±19 pA; I=0.08 mW/ μm^2^, n=9) (Fig.3E, Supplementary Fig. 3A) to saturation state at around 0.14 mW/μm^2^ (Fig. 3E, Supplementary Fig. 3A). In current-clamp, light-evoked depolarizations ranging from 10 to 27 mV were obtained in most of the cells with 0.07 and 0.12 mW/μm^2^ illumination (n=8) (Fig.3F, Supplementary Fig. 3B). Higher depolarizations (V=37 mV; I=0.07 mW/μm^2^, n=1) were measured in one cell, which exhibited particularly high photocurrents (445 pA; I=0.08 mW/μm^2^, n=1) (Fig.3F, Supplementary Fig. 4). Depolarization variability was likely due to differences in expression of the opsin and the variable presence of intrinsic voltage-gated ion channels. Previous studies have shown that visual stimulation could lead to a depolarization between 4 mV and 26 mV (15±7 mV Mean±SD) (Euler and Masland, 2000). These results suggested that the range of power that we used yielded physiological activations, i.e. comparable to the ones that would be evoked by visual stimulations.

**Figure 3.**
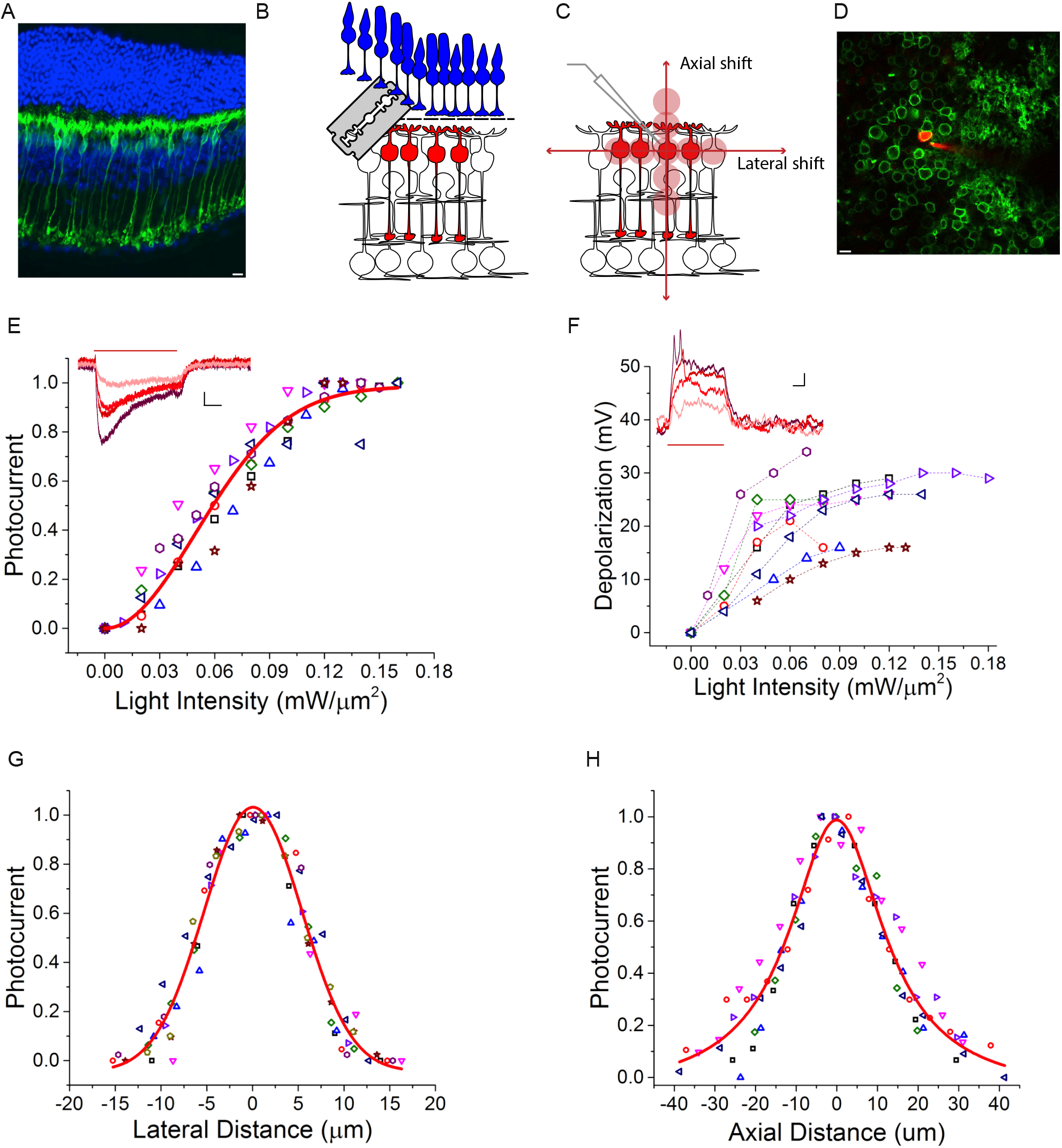
2P holographic stimulation enables physiological responses in RBCs with high spatial selectivity. **A:** Retinal slice showing the expression of CoChR-GFP in RBCs. Green: GFP. Blue: DAPI. Scalebar 10 μm. **B, C:** Schematic of the experiment. We first removed the photoreceptor layer by cutting the retina with a vibratome (see Methods), and then patched single RBCs expressing CoChR-GFP. We stimulated the cell with a 10-μm diameter holographic spot, which we successively moved laterally and axially to estimate the photostimulation selectivity. Power was set to depolarize the RBC in its physiological range. **D:** Representative two-photon image of a patched RBC. Green: CoChR-GFP-expressing cells. Red: Alexa594 dye, inside the pipette and filling the recorded cell. **E:** Peak photocurrent versus light intensity, normalized to the maximum for each recorded cell. Each symbol corresponds to a different cell. Red line: Saturation curve. *Inset:* Representative light-evoked photocurrents. Different traces correspond to different illumination intensities (from 0.02 mW/μm^2^ (light red) to 0.08 mW/μm^2^ (dark red) with 0.02 mW/μm^2^ steps). Vertical scale bar: 10 pA. Horizontal scale bar: 100 ms. Red horizontal bar indicates the photostimulation time (500 ms) **F:** Peak voltage depolarization in response to different stimulations, versus light intensity. Each symbol and dashed curve correspond to one cell. *Inset:* Representative light-evoked depolarizations. Different traces correspond to different illumination intensities (traces color code as in E). Vertical scale bar: 3 mV. Horizontal scale bar: 100 ms. **G:** Peak photocurrent as a function of lateral displacement, normalized to the maximum for each recorded cell. Each symbol corresponds to a different cell. Red curve: Gaussian fit. **H:** Same as G for axial displacements. Red curve: Lorentzian fit.

**Figure 4.**
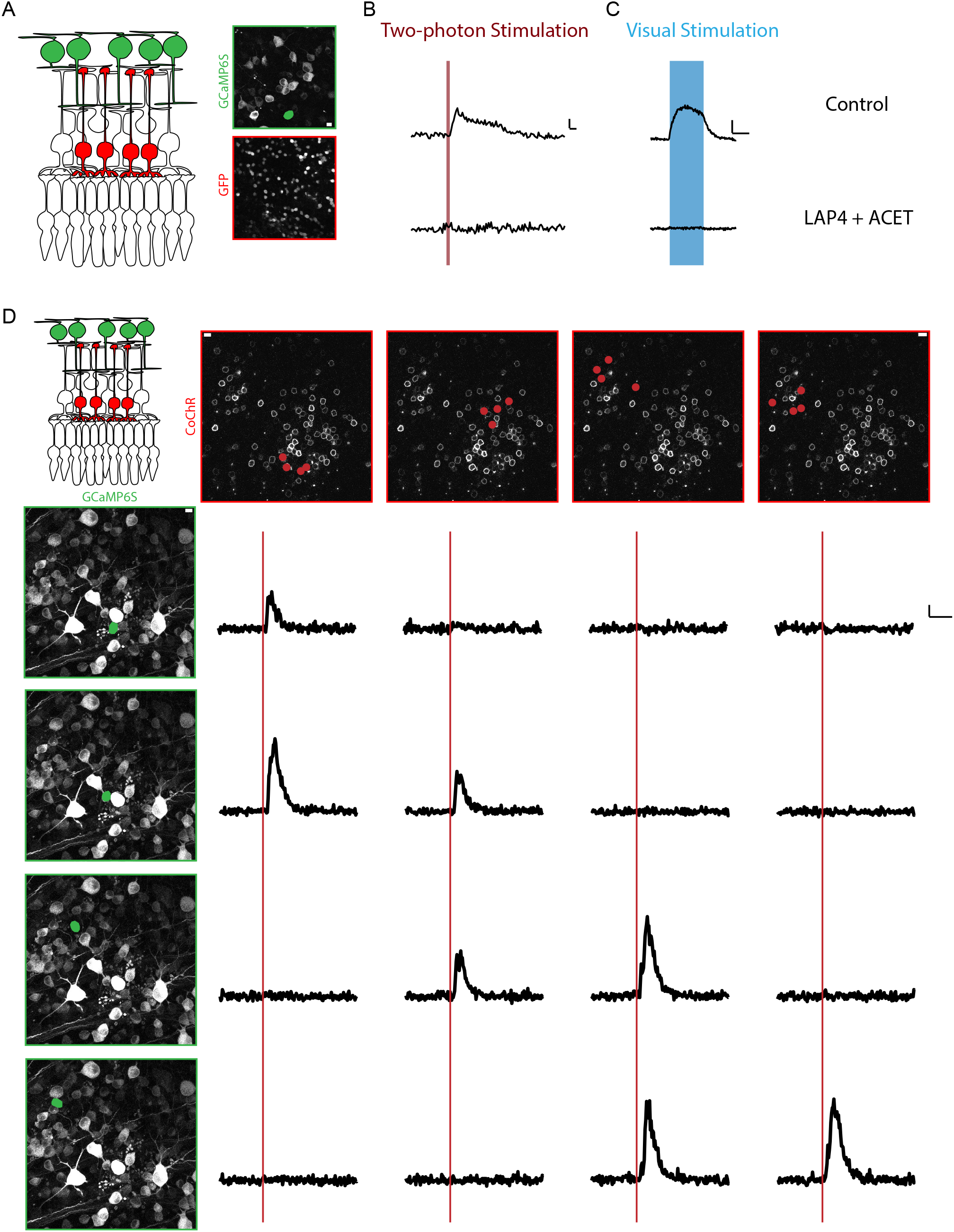
Patterned optogenetic stimulation of RBCs evokes selective activation of GCs. **A:** Control experiment. GCaMP6s is expressed in GCs, and GFP in RBCs. Top right: whole mount view of GCs expressing GCaMP6s. Bottom right: whole mount view of RBCs expressing GFP. **B:** Calcium response of a GC to a patterned holographic stimulation of RBC (500 ms, 0.1</<0.3 mW/μm^2^, indicated by a red line), under control conditions (top) and after bath application of LAP4 (20 μM) and ACET (1 μM) to block photoreceptor transmission to BCs (bottom). Scale bars: vertical DF/F 20%; horizontal 1 sec. **C:** same as B with visual stimulation with a flash of light (10 sec, P ~1,5 10^-3^ mW/μm^2^ indicated by the blue region). Scale bars: vertical ΔF/F 50%; horizontal 5 sec. **D:** All optical characterization of the retinal network, after LAP4+ACET application. GCaMP6s is expressed in GCs and CoChR-GFP in RBCs. Each column corresponds to a different pattern of stimulation in the RBC layer (red spots drawn in the top row). Each line corresponds to the response of a different GC (ROI indicated in green on the left column, on top of the GC layer image). Red line indicates the timing of the holographic stimulation (500 ms, 0.06<I<0.1 mW/μm^2^); Scale bars: 10 μm.

Next, we checked the spatial precision of the photostimulation spot. To this end we measured the photo-induced current while the holographic spot was laterally and axially displayed around the RBC soma (Fig. 3G), keeping the illumination intensity, I, in the physiological range (0.03 mW/μm^2^ < / < 0.09 mW/μm^2^). We observed a 50% drop of photocurrents by moving the spot 6μm laterally aside from the center of the cell (Fig. 3G). A residual 10% of photocurrent was recorded when the spot was 10 μm apart (corresponding to the lateral size of the illumination spot). Axially, photocurrents exhibited a decay of 50% at around 14 μm from the focal plane, with residual photocurrents below 10% for axial shifts superior to 30 μm (Fig.3H). Overall, the spatial selectivity of photoactivation was estimated to be 12 μm and 28 μm, corresponding to the lateral and axial FWHM of the photocurrents’ spatial distribution. These results show that photostimulation of RBCs could be achieved at cellular resolution.

### Simultaneous holographic stimulation of RBCs and imaging of GCs

Mice were co-injected with AAVs driving CoChR and GCaMP6s expression in the RBCs and GCs respectively (see Methods). We imaged the GC layer with 2P scanning imaging, while stimulating with one or multiple spots the RBCs, on average 70 μm deeper. In control retinas, where BCs only expressed GFP (Fig. 4A), we observed responses to holographic illumination (23/123 cells, n=2 retinas) that were due to out of focus photoreceptor stimulation (Euler et al., 2009)(Palczewska et al., 2014). To avoid photoreceptor stimulation and ensure calcium responses only evoked by holographic stimulation of RBCs, we blocked the transmission from photoreceptors to BCs by putting LAP4 and ACET, which block the transmission to ON and OFF BCs, respectively (see Methods)(Borghuis et al., 2014). Adding this pharmacological cocktail to the bath in control experiments abolished the responses induced by both holographic illumination (Fig. 4B) and flashes of light (44/84 cells, n= 2 retinas) (Fig. 4C).

All subsequent experiments were performed under pharmacological block allowing recording of calcium responses originating exclusively from holographic stimulation of CoChR expressing RBCs (see Methods).

Stimulation with one holographic spot rarely evoked any visible calcium response in GCs (0.15±0.07% of 5540 RBC-GC pairs tested, n=3 experiments) whereas simultaneous stimulation with *n* multiple spots (2 ≤ *n* ≤ 8) evoked reliable responses. Calcium responses were specific to the stimulation pattern: different calcium response patterns obtained in response to different BC stimulation patterns are shown in Figure 4D. As expected, the GCs closest to the stimulation pattern were the most likely to respond (Fig. 4D). Overall, the probability of responses decreased with distance.

### Functional asymmetry in a DS circuit

We next aimed to understand how DSGCs integrate the stimulation of RBCs. To detect DSGCs we first performed calcium imaging while stimulating photoreceptors with a full-field stimulation (Baden et al., 2016) and with bars moving in different directions to determine the tuning and polarity of the imaged ganglion cells (Fig. 5A). We then blocked the photoreceptor-BCs transmission as previously described and stimulated RBCs with multiple patterns (Fig. 5B,C).

**Figure 5.**
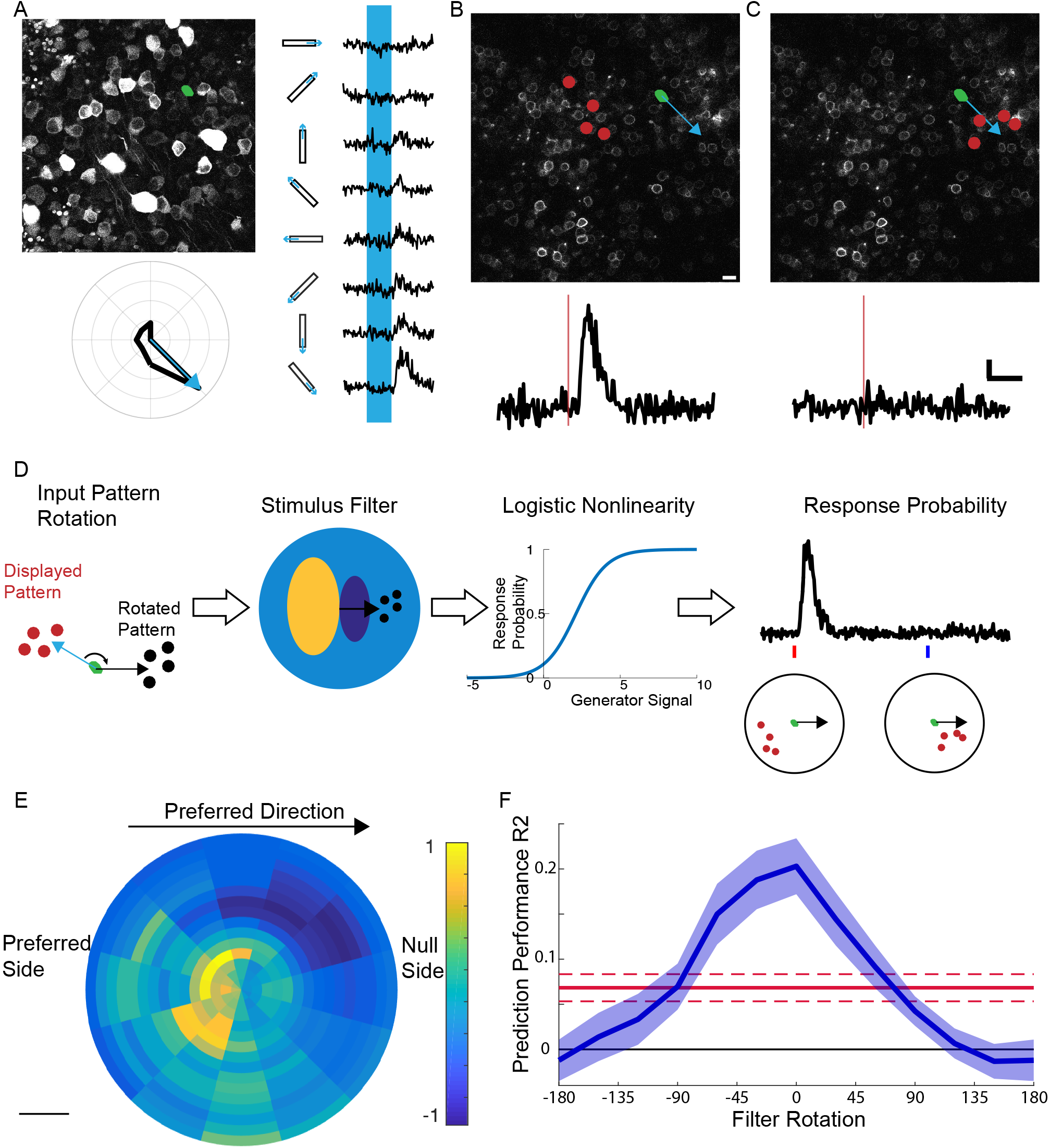
OFF DSGC integrate RBC output in an asymmetric manner. **A:** example of an OFF DSGC (G_2_ type). Top left: field of view in the GC layer, with GCaMP6s expressed in GCs. Green spot: ROI of one OFF DSGC. Right: calcium response of the same OFF DSGC cell to white bars moving in different directions. The blue rectangle indicates when the white bar is on the GC. Bottom left: polar plot showing the peak response of the same cell for each motion direction. Blue arrow: preferred direction (see Methods). **B:** *Top:* whole mount view of RBCs expressing CoChR, in the same retina as A. Red spots: holographic stimulation pattern. Green spot: corresponding location of the ROI of the OFF DSGC shown in A, with preferred direction shown with blue arrow. *Bottom:* calcium response of this GC to this holographic pattern. **C:** same as B for the same cell but a different pattern, located on the null side. **D:** schematic of the logistic regression analysis. Each stimulation pattern is first rotated to align all the preferred directions of all OFF DSGCs to the right. The model filters the stimulation pattern and the result goes through a sigmoid to predict the probability of response to each pattern. **E:** Filter estimated from the population of OFF DSGCs (G_2_) recorded. Scale bar: 50 μm. **F:** Prediction performance of the regression on a testing dataset (blue, R^2^-mean and s.e.m.) against the rotation angle (0: no rotation, see text). Red: Prediction performance under the hypothesis of a symmetric filter (see text, Mean ± SD).

Among the DSGCs, we focused on a specific OFF DS type described by Euler and colleagues as OFF DS G_2_ cell (Baden et al., 2016) (see Supplementary Fig. 5-note that these cells are different from the JAM-B cells described in (Kim et al., 2008)). For this cell type, we found that RBC stimulation pattern on the null side of the cell (“in front of the cell”, Fig. 1) did not evoke any response (Fig. 5C), while a similar pattern projected at an equal distance but in a different direction evoked strong responses (Fig. 5B). This suggests the possibility of asymmetric integration of RBCs output signal in OFF DS G_2_ cells.

To test if this bias applies at the population level, we performed a logistic regression test to each responsive region around the GC. This statistical model takes each pattern, aligns it to the GC’s position and preferred direction, and assigns a weight to each spot depending on its relative position. The sum of these weights is then converted into a probability to detect a calcium response. The weights are learned from the data, and form together a filter that tells which region is the most sensitive to stimulation. If the filter is asymmetric, it means that the RBC output is integrated asymmetrically. Note that this is equivalent to fitting a Linear-Nonlinear (LN) model (Chichilnisky, 2001) to the data.

To fit this model and learn its parameters, we first pooled together all the cells of this same OFF DSGC type, after realigning their position and direction preference (Fig. 5D). We fitted our model to these realigned data and measured its performance in predicting the responses of a test dataset not used for the learning. The model has a significantly better prediction performance (R^2^ = 0.2) compared to a null model (R^2^ =0.00), demonstrating that the filter inferred is meaningful.

The resulting filter was clearly asymmetric, biased against the preferred direction (Fig. 5E). We estimated its preferred angle to be 210 degrees (see Methods). To test if this asymmetry was significant, we rotated the filter and predicted again the responses to the stimulation patterns. Performance dropped to 0 for a filter oriented towards the null side (Fig 5F). It was larger for no rotation, but similar for a filter directed towards the preferred side. We randomly rotated each pattern before learning the model to ensure deviations from symmetry effects were not due to noise (see methods). The performance of the obtained shuffled models (Fig 5F) was significantly lower than our inferred model (p<10^-4^).

In the light of these results we can now choose between our two hypotheses. In the first hypothesis where RBC output is integrated asymmetrically, these GCs should respond more to patterns located in a specific direction, and our estimated filter should be significantly tuned for a specific direction. In the second hypothesis, the direction of the pattern has no influence on the cell response, and the filter should not be direction-selective. Our results suggest that the first hypothesis prevails, and the OFF G_2_ DS cells integrate RBC output asymmetrically.

## Discussion

### Multiplexed temporal focusing light shaping and functional imaging

We have established an all-optical method, for non-invasive interrogation of retinal circuits across axially distinct planes and have applied this approach to dissect how RBCs contribute to shape DSGCs.

Our optical system combines 2P multi-target temporally focused holographic illumination with a 2P scanning microscope, to independently activate one or multiple RBCs while monitoring the GCs response via 2P calcium scan imaging. The system also allows stimulation of the photoreceptor layer to classify the different cell types in the GC layer such as a specific type of DS ganglion cell: the OFF DS G_2_.

Multi target, multi plane, optical stimulation and functional imaging have been achieved with conventional (without TF) 3D-CGH in zebrafish using small holographic spots (6-μm diameter) (dal Maschio et al., 2017) and with TF low-NA Gaussian beams (Mardinly et al., 2018) or multi-spiral-scanning in mammals (Yang et al., 2018). CGH-only configurations are limited to small-size targets (dal Maschio et al., 2017), as radially enlarging the illumination pattern quickly deteriorates the axial confinement of the excitation (Bègue et al., 2013). On the contrary, in agreement with previous findings (Accanto et al., 2018; Hernandez et al., 2016) we have shown that 3D-TF-CGH enables a 2-3 fold improvement of the confinement allowing us to maintain the axial confinement independently from the lateral pattern extension. 3D multiplexing of TF low-NA Gaussian beams (Mardinly et al., 2018) enables multi target optical stimulation but the use of a rotating diffuser as opposed to the fixed holographic phase mask, used in our system, deteriorates axial resolution by 2-3 folds (Chen et al., 2018a; Pegard et al., 2018). Moreover, unlike the top-hat intensity profile of a holographic beam, the use of a Gaussian intensity profile can deteriorate the lateral resolution at powers close to saturation. Spiral scanning of multiple holographic spots enables multi target, multi plane stimulation (Yang et al., 2018), but to our experience requires higher excitation intensity to activate CoChR (Picot et al., 2018). Moreover, due to the out-of-focus light excitation when using intensities close to saturation, this approach also gives a worse axial resolution (Andrasfalvy et al., 2010). Here, we have used a static phase mask to generate the holographic disk on the TF grating. More flexibility to tune the spot shape and size could be achieved by replacing the mask with an SLM (Accanto et al., 2018), although this will add complexity to the optical system. In this study distribution of cells in the GC layer is relatively flat, thus enabling to restrict 2P calcium imaging to a single plane. For the investigation of more complex circuits the system can be integrated with multi plane imaging strategies based on e.g. divergence control or multiplexing of the imaging beam (Ji et al., 2016).

### Retinal circuit investigation

Previous studies estimated the projective field of BCs onto GCs using a combination of BC intracellular recordings and GC extracellular recordings using multi-electrode arrays (Asari and Meister, 2012, 2014; Baccus and Meister, 2002). This approach is limited as it does not allow to target a single BC type nor the simultaneous targeting of multiple cells lowering the overall yield of these experiments.

Our all-optical method allows unbiased recordings from all GC types, which allows us to identify the OFF DSGCs among the hundreds of cells recorded. MEA recordings do not record equally well from all cell types in the mouse retina. Targeted patch recordings would require a genetic strategy to label specifically the G_2_ OFF DSGCs studied here, and no such strategy exists so far. Our method allows high throughput stimulation of many RBCs, while imaging many GCs, which in turn enables us to detect the cellular receptive field for any type of GC.

### Asymmetry in the direction selective circuits

Our results suggest that the RBC pathway preserves asymmetry despite its strong divergence. This asymmetric input in turn contributes to direction selectivity. The secondary and ternary pathways are one synapse away from the cone circuitry and are also likely to preserve asymmetry. Taken together, these findings indicate that direction sensitivity is robust against changes in luminance in these direction selective cells because every circuit converging towards these cells is selective when interrogated alone. There seems to be no room for a pathway that would provide an unselective signal. This might be a general principle for the design of robust feature selectivity in neural circuits: all the circuits involved should be feature selective by themselves.

Our study also uncovered a specific asymmetry, with a preference for stimulation on the preferred side, rather than the null side. This bias results from the underlying circuit generating direction selectivity. As described in the introduction, a common circuit for many DSGCs, including ON-OFF DSGCs (Briggman et al., 2011; Fried et al., 2002; Wei et al., 2011), is based on asymmetric inhibition (Fig. 1A), where DSGCs receive more input from inhibitory ACs located on the null side. This inhibition suppresses the responses to a bar moving from the null side. As a consequence, stimulating on the preferred side evokes a stronger response than on the null side, where the resulting inhibition will cancel excitation. However, all DSGC types do not share this bias for the preferred side. In several other DSGC types direction selectivity is due to an asymmetric dendritic tree, oriented towards the null side (Kim et al., 2008; Trenholm et al., 2011). This results in the opposite bias, i.e. a stronger response for stimulation on the null side. The circuit underlying direction selectivity in G_2_ OFF DSGCs is still unclear. Our results suggest that it might be similar to the one at work for ON-OFF DSGCs: an asymmetric inhibition that induces the observed bias for the preferred side.

## Conclusions

In conclusion, here, we present an experimental approach combining advanced optical methods with *ad hoc* genetic strategies for optogenetic targeting allowing an all-optical interrogation of a retinal circuit. We investigated the primary pathway implicated in night vision, which relays the rod signal to the G_2_ OFF DSGCs through RBCs, and we have shown that this pathway preserves direction selectivity.

Our approach is extendable to other brain regions opening the way for a precise interrogation *in vitro* and *in vivo* (Chen et al., 2018b) of multi-layered circuits, for example to investigate how the information is transmitted across the multiple layers of the visual cortex (Wertz et al., 2015; Yang et al., 2016). Understanding this information transfer is a promising avenue to dissect complex neural circuits and understand the neural basis of computations.

## Methods

### Animals

All experiments were done in accordance with the National Institutes of Health Guide for Care and Use of Laboratory Animals. The protocol was approved by the Local Animal Ethics Committee of Paris 5 (CEEA 34) and conducted in accordance with Directive 2010/63/EU of the European Parliament. All mice used in this study were C57Bl6J mice (wild type) from Janvier Laboratories (Le Genest Saint Isle, France).

### AAV Production and injections

Recombinant AAVs were produced by the plasmid cotransfection method (Choi et al., 2007) and the resulting lysates were purified via iodixanol gradient ultracentrifugation as previously described. Briefly, 40% iodixanol fraction was concentrated and buffer exchanged using Amicon Ultra-15 Centrifugal Filter Units (Millipore, Molsheim, France). Vector stocks were then tittered for DNase-resistant vector genomes by real-time PCR relative to a standard(Choi et al., 2007).

For injection, animals were anesthetized with Isofluorane (Isoflurin 250 ml, Vetpharma Animal Health) inhalation and pupils were dilated. A 33-gauge needle was inserted into the eye to deliver the vector into the vitreous. 2 μl of vector solution was injected per eye, containing 1 μl of the vector delivering GCaMP6s (containing ~ 10^10^ vg) and 1 μl of the vector delivering either CoChR (containing ~ 10^10^ vg) or GFP (containing ~ 10^10^ vg).

For all experiments we used GCaMP6s (Chen et al., 2013) under the SNCG promoter(Chaffiol et al., 2017) to specifically target ganglion cells and we used AAV2 as viral vector. To express CoChR (Klapoetke et al., 2014; Shemesh et al., 2017), we used a recently published promoter (In4s-In3-200En-mGluR500P) (Lu et al., 2016), which has been proved to allow specific expression of optogenetic proteins in RBCs. To deliver it across the retinal layers we used 7m8 a genetic variant of AAV2 (Dalkara et al., 2013). Finally we used GFP only under the grm6 promoter (Macé et al., 2015) delivered with AAV2-7m8 to target BCs in control experiments. The injections were performed in 4-5 weeks old mice.

### Tissue preparation

For all experiments, we used female mice 4-8 weeks after the injection. Animals were dark adapted for at least 1h, then anesthetized with isofluorane (Isoflurin 250 ml, Vetpharma Animal Health) and killed by cervical dislocation. The eyes were enucleated and placed in AMES medium (Sigma-Aldrich, St Louis, MO; A1420), bubbled with 95% O_2_ and 5 % CO_2_ at room temperature. The eyes were dissected under dim red light (>645 nm) and the isolated retinas were flat mounted with GCs up and transferred to the recording chamber in the microscope. The retina was continuously perfused with bubbled Ames’ medium at a rate of 5-7 mL/min during experiments and temperature was maintained around 34 degrees.

### Experiment description and pharmacology

At the beginning of the experiments the flat mounted retina was placed under the microscope and left to rest for ~30 min in the dark. The first step of the experiment was to perform the visual stimulations (see below) to determine which cells were direction selective. Then, to block the photoreceptors (Borghuis et al., 2014), we added to bubbled AMES medium the KAR selective agonist ACET (1uM, catalog no 2728, Tocris bioscience) and the metabotropic glutamate receptor agonist LAP4 (20 μM, catalog no 0103, Tocris Bioscience). The retina was left to rest in the dark for ~ 30-45 min. Before starting to stimulate the RBCs expressing CoChR, we tested that the photoreceptor transmission to BCs was effectively blocked by doing visual stimulations on a central FOV of 100×100 μm^2^. The highest intensity of light used to stimulate photoreceptors was 1,53 x 10^-3^ mW/mm^2^. As shown in (Shemesh et al.,2017) (Supplementary Figure 4) this power is negligible compared to the one necessary to induce any activation of the opsin, which has small responses for ~2 mW/mm^2^. If no ganglion cell was responding to the visual stimulation, we proceeded with the holographic stimulation.

### Single cell electrophysiology

#### Tissue preparation

4 to 5 weeks old mice were injected with 1 or 1.5 μl volume of AAV2-7m8 carrying CoChR (~ 10^10^ vg) under a promoter specific for RBCs(In4s-In3-200En-mGluR500P) (Lu et al., 2016). 4 to 10 weeks after the injection, the animals were anesthetized with isoflurane and killed by cervical dislocation. Eyeballs were enucleated and dissected under white light. To have a better access to the BCs with the patch pipette, we removed the photoreceptor layer using a vibratome (Leica VT1200S slicer). This procedure was previously described in details (Clérin et al., 2014). Briefly, the dissected retina was transferred in the vibratome tank filled with bubbled Ames. The retina was placed photoreceptors down on a gelatin block in the center of the tank and the solution was removed to permit the sealing of the flat-mounted retina. Once the retina was sealed, the tank was filled with bubbling Ames again and the vibratome’s blade was lowered until the GCs level. A slice of ~ 80-90 μm was cut and transferred to the recording chamber under the microscope with GCs down. BCs were thus on the upper side without the photoreceptors on top of them, which made them more accessible to patch recordings (figure 2B, C).

#### Patch-clamp recordings

BCs layer was imaged with a 2P imaging system to select the region with cells expressing the opsin. BCs were visualized with an IR illumination, a water-immersion 40x objective (40x W APO NIR; Nikon), and an IR CCD (see “Optical system”) while approaching the cell with the patch pipette. Light was turned off soon after the whole-cell configuration was established. Patch-clamp electrodes were pulled from borosilicate glass capillaries (1.5mm outer diameter, 0.86mm internal diameter; Harvard apparatus) with a horizontal micropipette puller (P1000, Sutter Instruments). Pipettes were filled with the following solution (mM): 130 K-gluconate, 7 KCl, 4 MgATP, 0.3 mM Na-GTP, 10 Na-phosphocreatine, and 10 mM HEPES (pH adjusted to 7.28 with KOH; osmolarity 280 mOsm). Pipette resistance in the bath was 4.5-6 MΩ. An Ag/AgCl pellet was used as reference electrode in the recording chamber. Patched cells were loaded with Alexa 594 (Invitrogen) added to the pipette solution to reconstruct the morphology at the end of each experiment.

Data were acquired with a MultiClamp 700B amplifier (Molecular Devices), a National Instrument board, and the Neuromatic software (www.neuromatic.thinkrandom.com) running on IgorPro interface (Wavemetrics). Voltage and current clamp recordings were low-pass filtered at 6-10 kHz and sampled at 20-50 kHz. Cells were clamped at −40 mV. The cell resting membrane potential (Vm) was measured soon after achieving the whole-cell configuration (Vm=42±10 mV, from 24 cells). Series resistance (Rs) was determined and compensated from 70 to 80% with the MultiClamp software during acquisition (Rs=18±7 MΩ, from 24 cells). Cell membrane capacitance (Cm) was 3.8±0.8 pF (from 24 cells). Voltage values shown are not corrected for the liquid junction potential (estimated value: 15 mV).

To determine the axial and lateral resolution of the system we stimulated the cell with a holographic spot and moved it in steps of 2.5 μm laterally or 5 μm axially to estimate the photostimulation selectivity. We determined the peak photocurrent for increasing 2P light intensities and we normalized photocurrents to the maximum value for each recorded cell. Photocurrent saturation curve in Fig. 2E was given by empirically fit data with (1 — *e-^x2/k^)* with k equals to 0.06.

### Optical System

The optical system was built around a commercial upright microscope (SliceScope; Scientifica) and combined a multi-light-path imaging architecture, a 3D multiplexing temporally focused holographic-based photoactivation apparatus and a spatiotemporally-controlled visual stimulation system.

The imaging system has been already described in (Ronzitti et al., 2017). Briefly, it includes three different imaging pathways: a 2P raster scanning, a 1P wide-field epifluorescence, and a wide-field infrared (IR) illumination imaging. 2P imaging was provided by a femtosecond pulsed beam (Coherent Chameleon Vision II, pulse width 140 fs, tuning range 680-1080 nm), relayed on a pair of XY galvanometric mirrors (3 mm aperture, 6215H series; Cambridge Technology), imaged at the back aperture of the microscope objective (40x W APO NIR; Nikon) through an afocal telescope. Galvanometric mirrors were driven by two servo drivers (MicroMax series 671; Cambridge Technology) controlled via a digital/analog converter board (PCI-6110; National Instrument) through ScanImage software (Pologruto et al., 2003). Emitted fluorescence was collected by two photomultiplier tubes (PMT) GaAsP (H10770-40 SEL; Hamamatsu #H10770-40 SEL) coupled to the objective back aperture via a fiber-coupled detection scheme (Ronzitti et al., 2017). 2P imaging laser power was tuned by combining an electrically controlled liquid crystal variable phase retarder (Meadowlark Optics #LRC-200-IR1) and a polarizer cube (Meadowlark Optics #BB-050-IR1). For image acquisition, we used ScanImage synchronized with the visual stimulation or CGH-excitation with a custom-made software running in MATLAB. For visual stimulation acquisitions, we divided a 200umx200um FOV in 4 100×100 μm^2^ smaller FOV and we took 64×64 pixel image sequences at 7.8 frames per sec (Imaging power (P) < 7 mW after the objective for all recordings). For image acquisitions during optogenetic stimulation, we recorded a 200×200 μm^2^ FOV in a 128×128 pixel image sequences at 5.92 frames per sec (P ranging from 9 mW to 15 mW). For high resolution morphology scans, we took 512×512 pixel images. 1P widefield imaging was provided by a LED source (Thorlabs #M470L2). 1P emitted fluorescence was collected through a tube lens (f= 200 mm), on a charge-coupled device (CCD) camera (Hamamatsu Orca-05G) after passing through a dichroic mirror (Semrock #FF510-Di02) and a visible bandwidth filter (Semrock FF01-609/181). 1P-and 2P-emitted fluorescence was separated through a movable dichroic mirror (70×50mm custom size; Semrock #FF705-Di01) and an upstream dichroic mirror (Chroma #ZT670rdc-xxrxt). IR illumination was provided by a custom-made external IR stalk lamp fitted near the microscope. IR light reflected by the sample was collected with an IR CCD (DAGE-MIT IR-1000). 2P optogenetic photoactivation was performed by generating 10 μm diameter circular spots pinpointing opsin-tagged cells in the sample via a 3D multiplexed spatially controlled phase modulation of the illumination beam wavefront thoroughly detailed in (Accanto et al., 2018). Specifically, a femtosecond pulsed beam delivered by a diode pumped, fiber amplifier system (Satsuma HP, Amplitude Systemes; pulse width 250 fs, tunable repetition rate 500-2000 kHz, gated from single shot up to 2000 kHz with an external modulator, maximum pulse energy 20 μJ, maximum average power 10 W, wavelength λ=1030 nm) operated at 500 kHz, was widened through an expanding telescope (x5) and transmitted through a custom-designed 5×5 mm^2^ 8-grey-levels static phase-mask calculated via Gerbergh and Saxton algorithm and fabricated by etching of fused silica (Double Helix Optics, LLC). The phase mask profile was expanded 2x and was Fourier transformed by a 500mm lens to project the holographic spot on a blazed reflective diffraction grating (600 l/mm) for temporal focusing.

The beam was then collimated through a 1000mm lens to impinge the sensitive area of a reconfigurable liquid crystal SLM (LCOS-SLM, X13138-07,Hamamatsu Photonics) placed in the Fourier plane of the diffraction grating. A beam stop was placed to physically block the SLM’s not modulated zero order. The SLM plane was imaged through a telescope on the back focal plane of the objective lens and addressed with a phase modulation calculated with a custom-designed software (Wavefront-Designer IV) to produce a set of diffraction-limited spots able to multiplex the circular spot in 3D at the sample plane and light-target opsin-expressing cells in the bipolar cells layer. For all experiments in Fig. 4 and 5, we used photostimulation intensities ranging from 0,06 mW/μm^2^ to 0,1 mW/μm^2^.

Visual stimulation was performed by spatiotemporally-controlled full-field or moving-bars visual stimuli generated through a DMD-based amplitude modulation. A 420nm LED beam (Thorlabs #M420L2) was filtered by a bandwidth excitation filter (Semrock FF01-420/10), conveniently attenuated with density filters and collimated to illuminate the sensitive area of a DMD (Vialux GmbH). The DMD plane was conjugated to the sample plane by a telescope through the rear port of the microscope. Visual stimuli were generated by a Matlab custom-designed software and synchronized with the 2P raster scan retrace. The LED intensity was calibrated to range (as photoisomerization rate, 10^10^ P */sec cone) from 0,3-2 (photoisomerization rate) and 1-5 to 39-43 and 120-130 for S and M opsins respectively. For all experiments, the retina was kept at constant intensity level for 30 seconds from the laser scanning start to the beginning of the visual stimuli. We used two types of visual stimuli: 1-full field ‘chirp’ stimulus (Baden et al., 2016) consisting of a bright step of 10 seconds and two sinusoidal intensity modulations, one with increasing frequency and one with increasing contrast; 2-0.3 x 4 mm bright bar moving at 1 mm s^-1^ in eight directions on a dark background.

### Data Analysis

Data analysis was performed using MATLAB. Region of interest (ROIs), corresponding to somata in the RGC layer, were identified semi-automatically using a custom software based on a high resolution image of the ganglion cell layer and on a projection of all the images acquired for each stimulation. Electrophysiological recordings were analysed with IgorPro (Wavemetrics) and OriginPro (OriginLab).

### Direction Selective Cells identification

#### Pre-processing

The Ca^2+^ traces for each ROI were extracted as (*F* − *F*_0_) /*F*_0_, where *F* is the mean fluorescence trace over the ROI, and *F*_0_ is the average fluorescence over the 5 seconds preceding the visual input. For each bar direction, we computed the median response *r_d_ (t)* across repetitions (three to six repetitions). Each median response was then normalized such that *ma x_d_*(*ma x_t_*(|*r*_*d*_(*t*)|)) = 1, for *d*from 1 to 8 directions and *t* running over the entire trace.

### Response quality index

To measure how well a cell responded to a stimulus, we computed the signal-to-noise ratio as in (Baden et al., 2016):

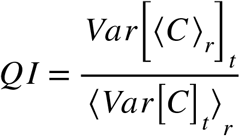

Here *C* is the response matrix from time samples *T* by stimulus repetitions *R*. Each row in *C* is the concatenation of the responses to all the 8 directions and each column is one repetition. ⟨ ⟩*_x_* and *Var* [ ]*_x_* are respectively the mean and the variance on the x dimension. *QI* is a global measure of the consistency of the responses to the moving bars.

We set the *QI* threshold to 0.2, meaning that each trace with *QI* below this value was discarded and not considered for further analysis.

### Direction and orientation selectivity

To extract time course and directional tuning of the Calcium response to the moving bar stimulus, we performed an analysis similar to the one described in (Baden et al., 2016). Briefly, we first performed a singular value decomposition (SVD) on the response matrix composed of the average response to each direction. We define the tuning curve V*(ϑ)* as the direction-dependent component of the first singular value. To measure direction selectivity (DS), we then projected the tuning curve on a complex exponential *φ_k_* = *e^iα_k_^* where *α_k_* is the direction in the *k*-th condition:

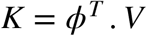

We computed a *DS* index as the resulting vector length

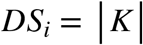

We labeled as *direction selective* each cell whose *DSi* value exceeded a given threshold (*DS_T_* = 0.7).

### Logistic regression

To fit our model, we first binned each stimulation pattern by dividing the space around each ganglion cell in 15 different radii from 0 to 150 μm, and in 12 different angles of equal size. Each stimulation pattern it was thus transformed into a vector *S_i_*, which our model used to predict the probability of response *p_i_*:

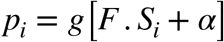

where *F* is the model filter, *g* the sigmoid function, and *α* a constant. This is the model used for logistic regression, and this model is also analog to a LN model (Chichilnisky, 2001) with a sigmoid as non-linear function and no temporal integration.

We learned the parameters *F* and *α* with a leave one out strategy, where all but one stimulation patterns are used as a training set and the remaining one is used for testing, and we iterate over all stimulation patterns.

We then maximized log-likelihood with a *L1* penalty for sparseness of the parameters, and a *L2* smoothness constraint between neighbouring values of the filter. The weights of these two cost functions was chosen such that the log-likelihood of the testing set was maximal.

The performance of the model was evaluated using the ***R***^2^ introduced by Tjur for logistic regressions (Tjur, 2009):

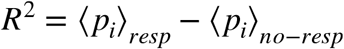

Statistical error on ***R***^2^ have been computed as error of the mean.

In order to compute the preferred direction of the inferred filter, we first averaged it over the radial coordinate and then applied the same projection strategy used before for the estimation of the cell preferred direction.

To test the hypothesis of an isotropic filter, we learned the model parameters with the same strategy, except that we rotated each stimulation with a random angle before learning. We repeated this random rotation many times to estimate the mean and standard deviation of the performance in this isotropic hypothesis.

