## Supplementary figures for "All-optical interrogation of a direction selective retinal circuit by holographic wave front shaping"

### Supplemental Information:

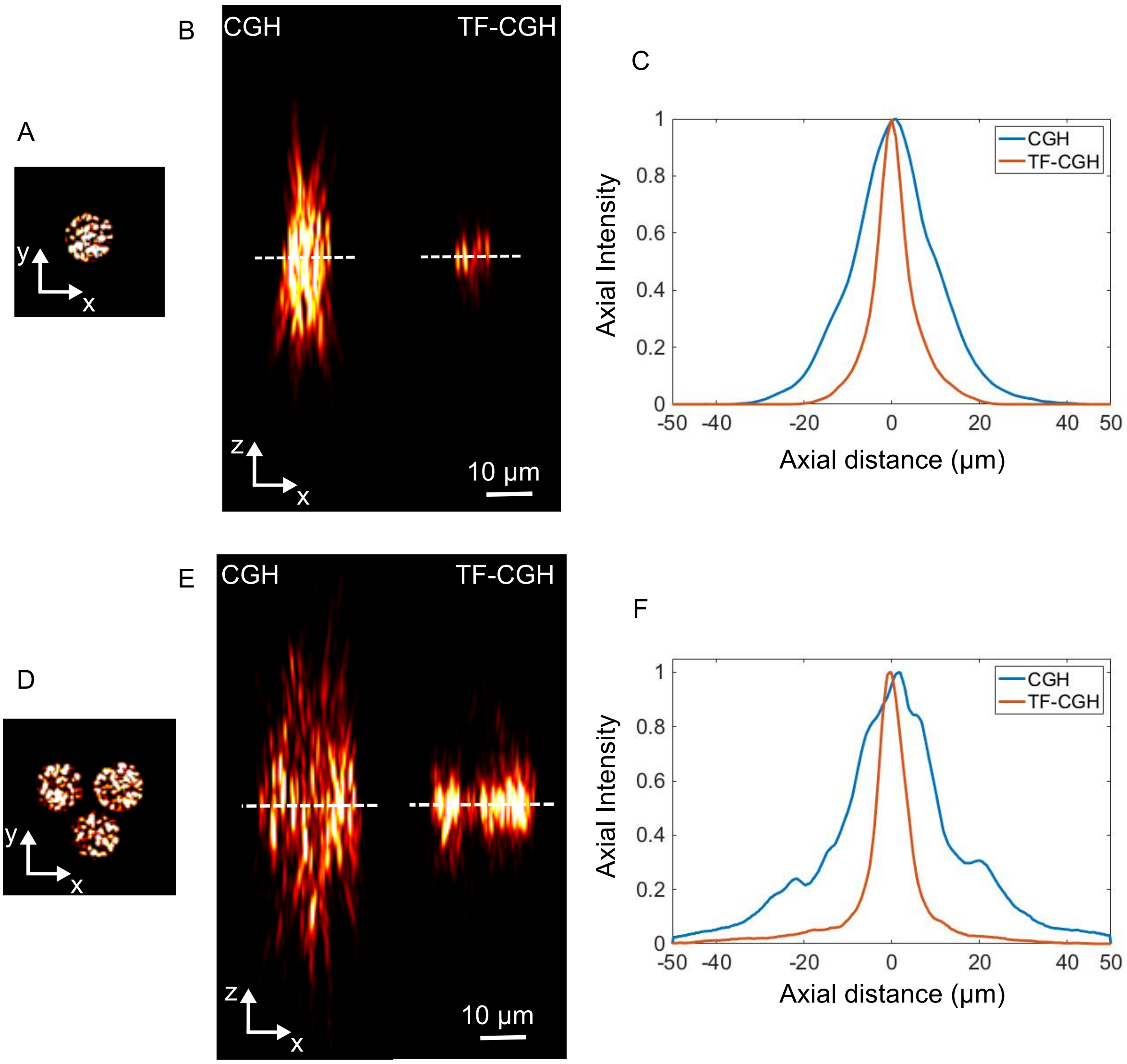


**Supplemental Figure 1. Comparison of axial resolution of temporally and non-temporally focused holographic spots. A.** Image of a 10-μm-diameter holographic spot. **B.** Simulation of the axial propagation of the spot in A without (left; CGH) and with temporal focusing (right; TF-CGH). The simulation spans in ±50 μm around the focal plane (dashed line). The images show the *xz* orthogonal maximum simulated fluorescence intensity projection of the spots. **C.** Axial profile of the simulated fluorescence intensity of spots shown in B for the two cases of CGH (blue line) and TF-CGH (red line). The FWHM of the axial profile is 17 μm for the CGH spot, and 7 μm for the TF-CGH spot. **D.** Image of three 10-μm-diameter holographic spots placed at a distance of 12.5 μm one from the other (center to center) arranged in a triangle-configuration. **E.** Simulation of the axial propagation of the spots in D without (left; CGH) and with temporal focusing (right; TF-CGH). The simulation spans in ±50 μm around the focal plane (dashed line). The images show the *xz* orthogonal maximum simulated fluorescence intensity projection for the excitation pattern of three spots. **F.** Axial profile of the simulated fluorescence intensity of spots shown in E for the two cases of CGH (blue line) and TF-CGH (red line). The axial profiles are plotted by integrating the simulated fluorescence intensity in an area covering the spots. The FWHM of the axial profile is 20 μm for the CGH spots, and 7 μm for the TF-CGH spots.

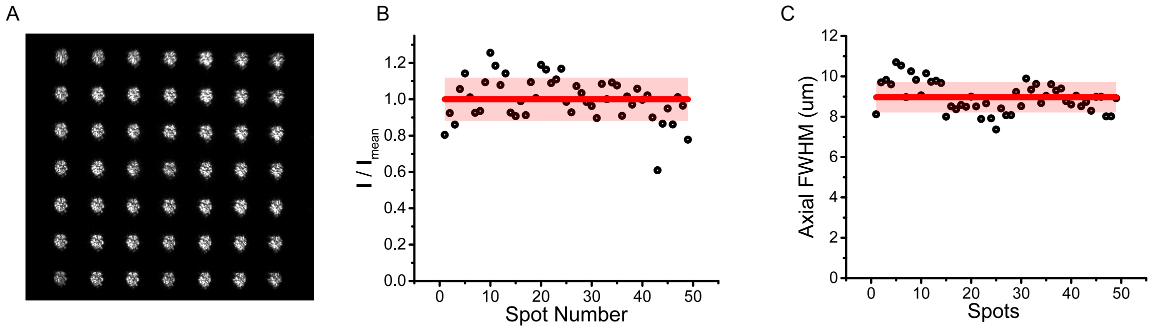


**Supplemental Figure 2. The optical system gives homogeneous photostimulation across the field of view**. **A.** Set of 10 μm diameter holographic spots simultaneously displayed 70 μm below the focal plane across the field of view. **B.** Normalized intensity of the different spots displayed in A. **C.** Axial FWHM of fluorescent intensity profile induced on a Rhodamine-6G layer by a set of holographic spots disposed as in A. Red solid line and light red band in B and C correspond to mean and standard deviation, respectively.

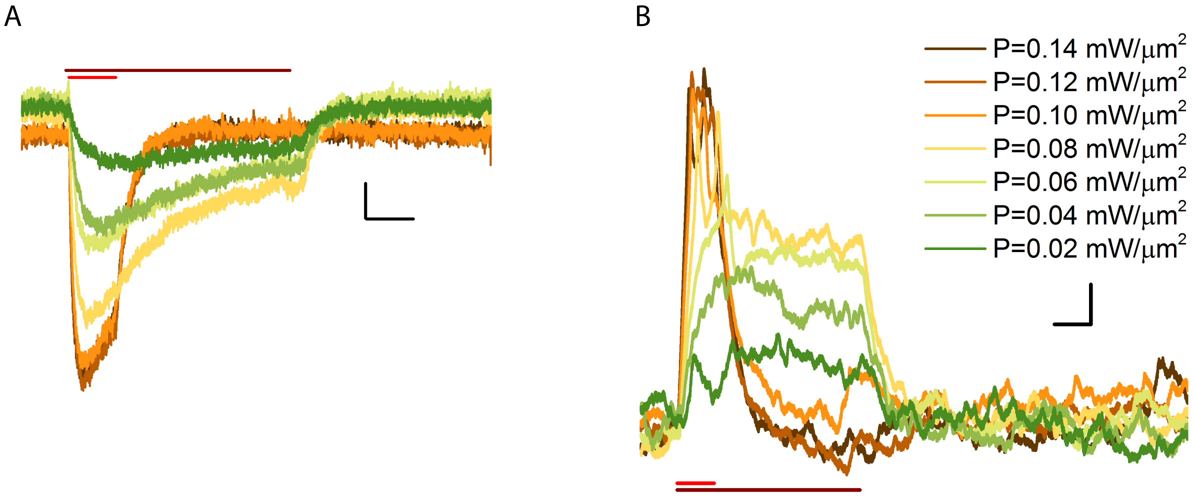


**Supplemental Figure 3. 2P holographic photo-stimulation enables physiological responses in RBCs**. **A.** Representative light-evoked currents and membrane depolarization **(B)** induced on a CoChR-expressing RBC under 2P holographic illumination at different powers. Red horizontal bars indicate the illumination period (500 ms for powers from 0.02 mW/µm$^2$ to 0.08 mW/µm$^2$; 100 ms for powers from 0.10 mW/µm$^2$ to 0.14 mW/µm$^2$). Vertical scale bar is 10 pA in A and 3 mV in B. Horizontal scale bar is 100 ms.

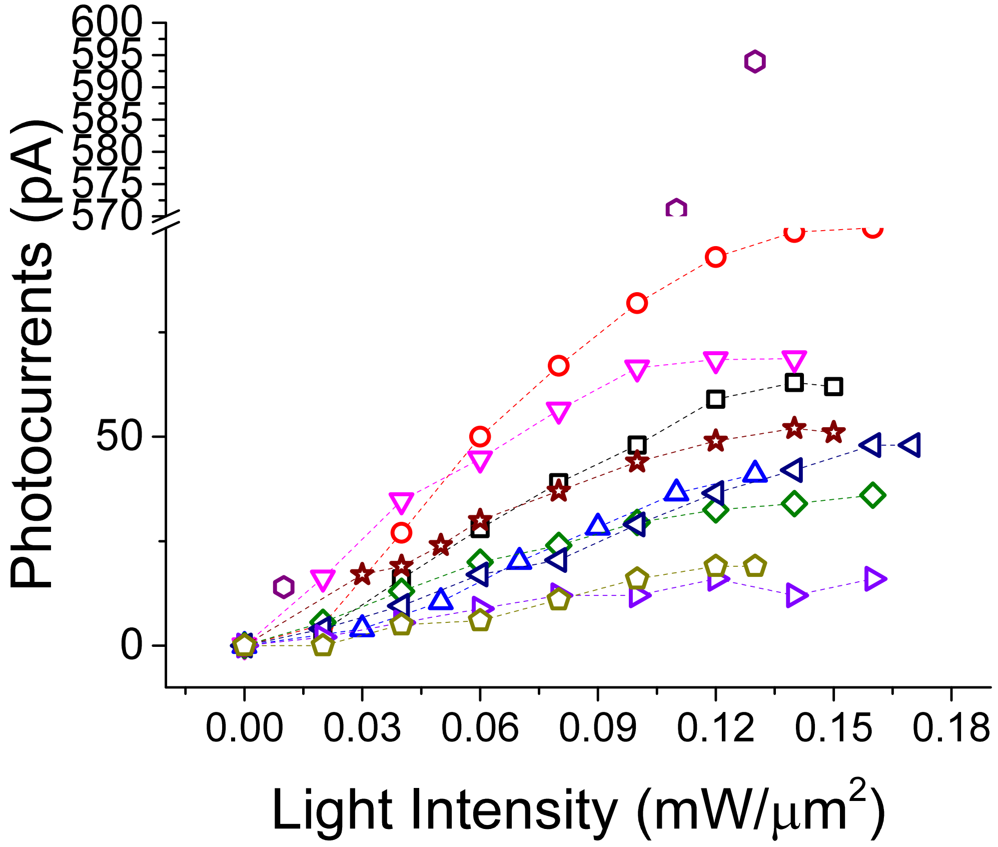


**Supplemental Figure 4. 2P holographic stimulation induces photocurrents of tens of pA.** Peak of the light-evoked current induced by photostimulating CoChR-expressing RBC under 2P holographic illumination at different powers. Different symbols indicate different cells.

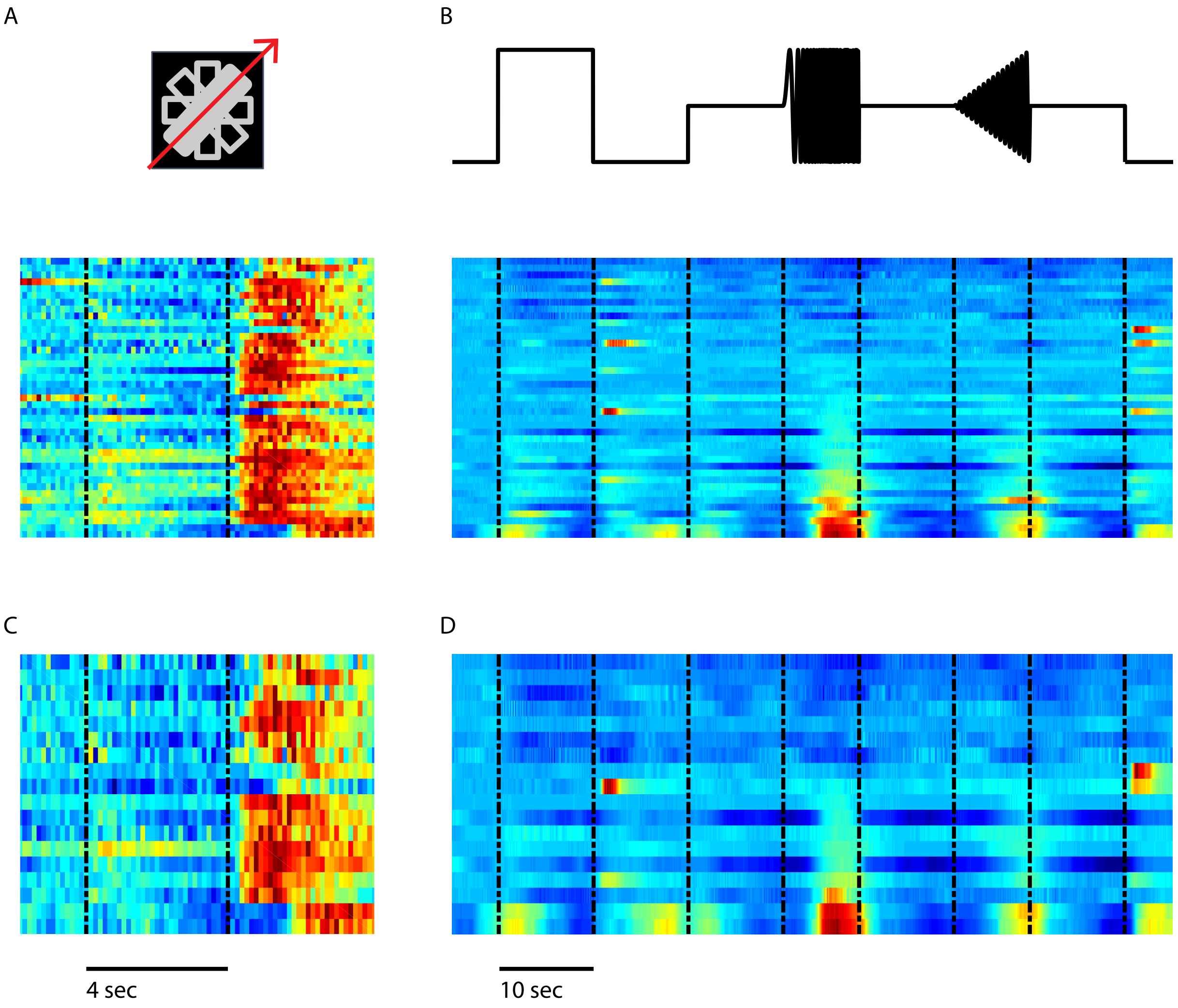


**Supplemental Figure 5. Identification of $G_2$ OFF DSGC.** We selected our subset of OFF DSGCs on the basis of their response to the moving bar: they had large OFF responses and no or little ON responses to the moving bar **(A, C)**. We also displayed a full field stimulus similar to the one used in (Baden et al., 2016) and observed responses similar to the ones observed for the $G_2$ type **(B, D)**: a response to the OFF flash and to the 'chirp' stimulus. Note that, similar to (Baden et al., 2016), we observed that some cells had little to no response to this full field stimulus, which might be caused by a strong surround inhibition. Heat maps illustrate single responses (max = 1). In panel A, B all the cells of the cluster are shown. In panel C, D only the cells responding additionally to the holographic stimulation are shown.
